# Evaluation of Bilateral Asymmetry in the Humerus of Human Skeletal Specimen

**DOI:** 10.1101/694984

**Authors:** Samuel S. Dare, Godfery Masilili, Kintu Mugagga, Peter E. Ekanem

**Affiliations:** Department of Human Anatomy, Faculty of Biomedical Sciences, Kampala International University Western Campus, Uganda; Anatomy Unit, Kabale University School of Medicine, Kabale University, Uganda; Department of Human Anatomy, School of Health Sciences, Makarere University, Uganda; Anatomy Unit, Biomedical Department, Mekelle University, Ethiopia

**Keywords:** Humerus, Bilateral asymmetry, humeral torsion, osteological, skeleton

## Abstract

Several studies have established a relationship between morphological and behavioural asymmetry making investigations of bilateral bone asymmetry an attractive and important research area. The purpose of this study was to investigate bilateral asymmetry patterns of skeletal specimen from five geographical locations (Rwanda, Burundi, Congo, Kenya and Uganda) at Galloway Osteological Collection, Department of Anatomy, School of Biomedical Sciences, Makerere University College of Health Sciences at Makerere University.

The angle of torsion and retroversion, mid shaft circumference, length and weight of 232 pairs of humeri were determined. A Torsiometer was used to measure the angle of torsion in degrees according to Krahl and Evans 1945, a tape was used to measure the mid shaft circumference at the level of the apex of the deltoid V and the length in cm was determined. An osteomeric board was used to measure the length of the humerus in centimeters. A weighing balance was used to measure the weight of the humerus in grams.

The analysis of humeral asymmetry with respect to parameters of the human skeletal specimen at the Galloway Osteological collection Mulago revealed bilateral asymmetrical status observed in the angle of torsion, length, weight and mid-shaft circumference. Our result mostly showed lateralization to the right in all the parameters investigated except the torsion angle which is to the left.

Our investigation revealed that humeral torsion is inversely proportional to weight, length and mid-shaft circumference of the humerus. This study established the existence of bilateral asymmetries in the humeri of all the geographical regions investigated with more asymmetry observed in the male compared with the female.

## INTRODUCTION

Symmetry is defined as correspondence in size, shape and relative position of parts on opposite sides of a dividing line or median plane while asymmetry is described as a lack or absence of symmetry. Although bilateral symmetry in paired morphological traits is evident in humans, however significant deviation from this observed in internal organs, human brain and especially in upper limb is referred to as bilateral asymmetry (Zaidi 2011).

Several studies have established a relationship between morphological and behavioural asymmetry making investigations of bilateral bone asymmetry an attractive and important research area (Steele 2000; Cuk*et al.* 2001; Lazenby 2002). This field may help to understand how behavior can influence the dynamic development of bone structure. Bone asymmetry is thought to basically results from disproportionate mechanical stress which influences bone remodeling and plasticity (Trinkaus *et al.* 1994; Churchill and Formicola 1997). Krahl*et al.* 1994; Bass *et al.* 2002 and Kontulainen *et al.* 2002 observations of asymmetry between playing and non-playing arms of tennis players revealed a strong effect of behavioural use of the limbs on diaphyseal structure.

According to Ruff, 2000, it is reasonable to believe that more active humans characterized by activities that are greatly influenced by mechanical stressors in life demonstrate greater asymmetry. Though genetic constitution may play some little role but external factors are believed to be major determinant of bilateral asymmetry. Studies have reported sex differences in bilateral asymmetry but results are not consistent depending on the skeletal sample and element (Ruff and Jones, 1981; Steele, 2000)

Asymmetry between the upper limbs bones have been reported in previous studies with little differences in all races and difference significantly greater in males than females (Warren, 1897; Schultz, 1937; Hiramoto, 1993; White & Folken, 2005). In most cases the left bones have been reported to be more variable in weight and length but the average lateral asymmetry was to the right in the arms. According to Latimer and Lowrance (1965) in the study of the weights and lengths of right and left bones of each pair from 105 human skeletons from Asia reported that all of the long bones of the upper limb were heavier and longer on the right side and that the humerus was most asymmetric.

The examination of the upper and lower limb asymmetries can be useful to medical scientists, archeologists, and anthropologists (Iscan & Shihai 1995; King *et al.*, 1998), to the police and forensic experts and for medicolegal studies (Steyn & Iscan 1999; Mall *et al*. 2001). Significantly, this intra-individual variation in the size and shape of the left and sides of the body has been linked with low back pain (Friberg, 1983; All-Eisa*et al*, 2004). All-Eisa*et al* (2004) showed that the higher the degree of asymmetry in the upper and lower limbs, the greater the likelihood of low back pain.

The consistent use of a limb habitually over the other during bimanual activities without a pathological condition result into asymmetric increase in the mechanical load of the preferred hand and limb dominance (handedness) (Roy *et al.* 1994; Cassandra, 2012). Studies on past populations which examined paired elements of the human skeleton for bilateral asymmetry thought it to be sexually dimorphic and associated handedness with dominant upper (Jaskulska 2009).

Past studies have established a relationship between hand dominance and side of asymmetric disease with right-handed individuals prone to more serious disease on the right side of the body. Body parts asymmetry such as forehead, facial structures, cubital and popliteal crease and thumbs has been found to be common in patients with localization related epilepsy syndromes.

Fong *et al.*, 2003 study of patients with seizure disorders concluded that body asymmetries is a useful clue to diagnosis of localization related seizure and may provide clues for lateralizing seizure origin in partial onset seizures.

Singh (1979) showed bilateral asymmetry in the direction and degree of tortion in metacarpal bones. According to Whiteley, (2009), “The development of humeral torsion seems to be determined by both hereditary and activity related factors (Edelson 2000; Krahl 1947) with the relative contributions of each remaining unknown. It is hypothesized that for the majority of adults, humeral torsion is largely genetic in origin, with opportunity available for activity-related influences of a less magnitude.

The purpose of this study was to investigate bilateral asymmetry patterns of skeletal specimen from five geographical locations (Rwanda, Burundi, Congo, Kenya and Uganda) at Galloway Osteological Collection, Department of Anatomy, School of Biomedical Sciences, Makerere University College of Health Sciences at Makerere University.

## MATERIALS AND METHODS

This study was done at Galloway Osteological Collection, Department of Anatomy, School of Biomedical Sciences, Makerere University College of Health Sciences at Makerere University. It consists of over 232 sets of skeletons both female and male and of different ages and geographical origin. It was cross-sectional descriptive and quantitative study involving measurements.

### Determination of humeral weight

A weighing balance with a margin of error of +/− 0.1 was used to measure the weight of the humerus in grams (g). Fig. 1

**Fig. 1:**
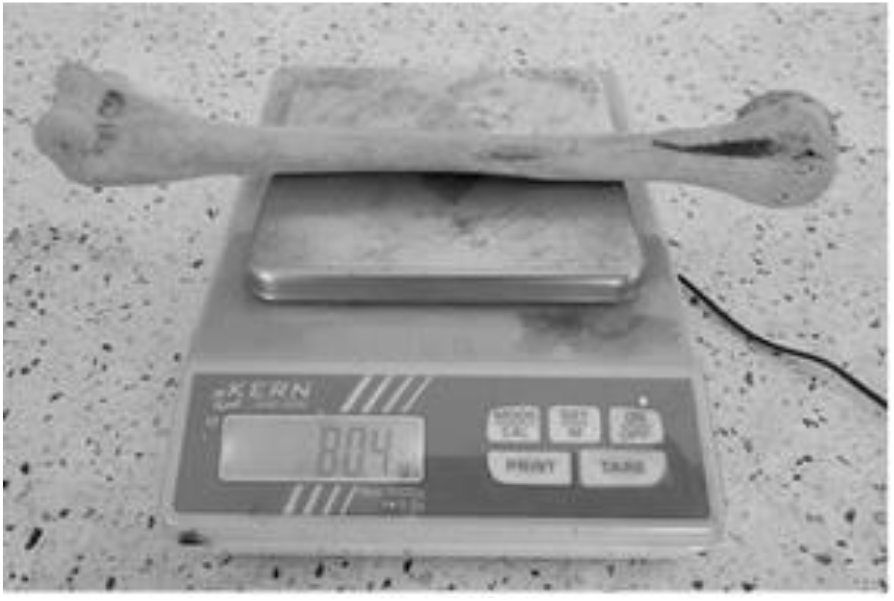
Measurement of humeral weight using a digital (KERN 440 −45N) machine

### Determination of Humeral length

An osteomeric board with a narrow margin of error of +/− 0.1 was used to measure the length of the humerus in centimeters (cm) by placing the bone horizontally on the board. Fig. 2.

**Fig 2:**
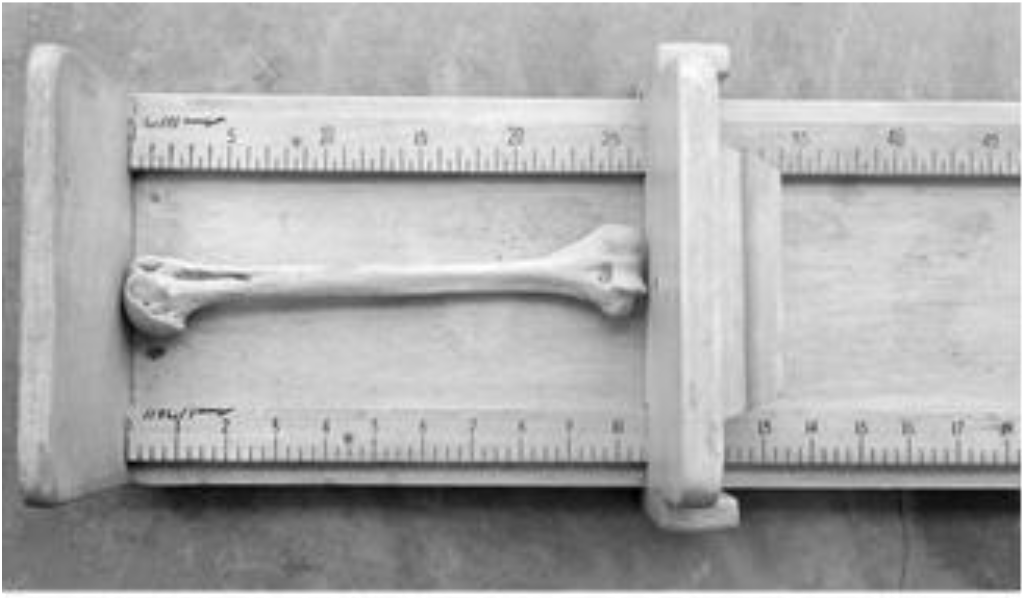
Measurement of humeral length using an osteomeric board

### Determination of Mid-shaft circumference

A millimeter graph paper was used to measure the mid shaft circumference at the level of the apex of the deltoid V in centimeters (cm). Fig. 3.

**Fig. 3:**
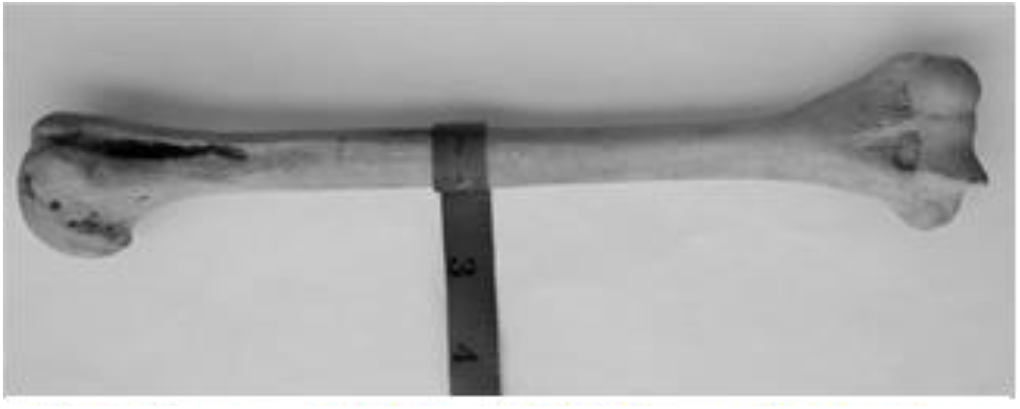
Measurement of humeral Mid-shaft circumference using a measuring tape

### Determination of angle of Torsion

A Torsiometer was used to measure the angle of torsion in degrees according to Dare et al, 2012. Fig. 4.

**Fig. 4:**
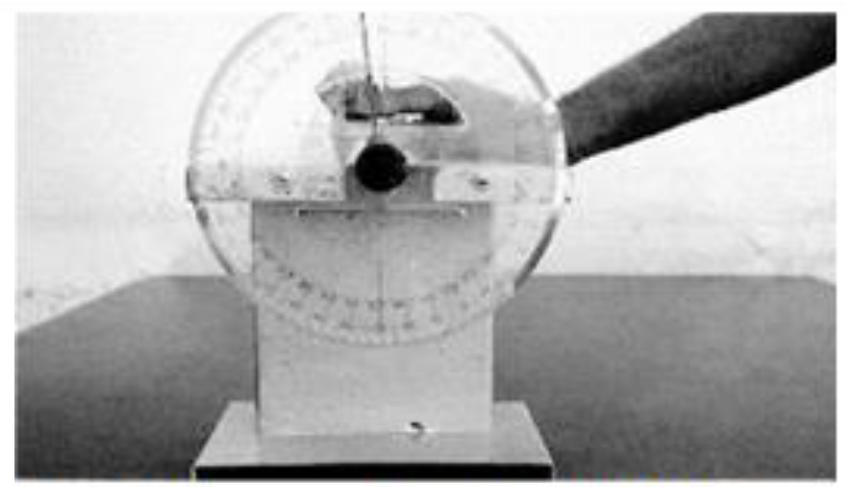
Measurement of humeral angle of torsion using a Torsiometer.

## DATA ANALYSIS

Data collected was analyzed scientifically using GraphPad Prism 7. All data was expressed as Mean +/− SEM. The data was analyzed using correlation coefficient and paired sample t-test with multiple comparisons. A P-value of <0.05 is considered as significant.

## RESULTS

The analysis of humeral asymmetry with respect to parameters of the human skeletal specimen at the Gallow Osteological collection Mulago revealed bilateral asymmetrical status observed in the angle of torsion, length, weight and mid-shaft circumference. The coefficient of correlation between parameters and p values were expressed as ‘r’ and ‘p’ respectively.

## DISCUSSION

Differences have been observed within or between body structures such as the size and shape of limb bones. According to (Roy *et al.* 1994; Ercan *et al*., 2008; Schaefer *et al.*, 2006) healthy individuals presents mild directional asymmetry which may result for example, from increased mechanical stress of the preferred limb over the other during habitual activities. Several parameters of the body long bones such as weight and length may reveal the degree of asymmetry. Steel and Mays (1995), measured the maximal length of the humerus, radius and ulnar in a series of 271 skeletons from medieval osteologic collection and reported the presence of oriented asymmetry in the arm bone length.

In our study, we measured the length, weight, torsion angle and mid-shaft circumference of the humerus in order to investigate the presence of bilateral asymmetry in different geographical locations. Our result shows the male and female right humerus heavier than the left in all the geographical locations except in the Congo and Kenya female humeral where the left was heavier than the right (Fig. 5). This is consistent with previous studies which reported that the long bones of the upper limb are heavier on the right side (Latimer & Lowrance 1965; Gutnik er al., 2015). The disparity observed in Congo and Kenya female may be due to fewer sample size and therefore cannot be conclusive. In figure 6, showing the comparison male and female corresponding right and left humeral weight, our result revealed a heavier right humeri in the male in all the geographical locations except in Kenya female which has a heavier humerus on the left compared to the left. This right sided asymmetry in humeral weight may be attributed to more frequent use of the right arm resulting in heavier or stronger muscles of that side and consequently heavier and stronger bones (Kewal 2011).

**Figure 5:**
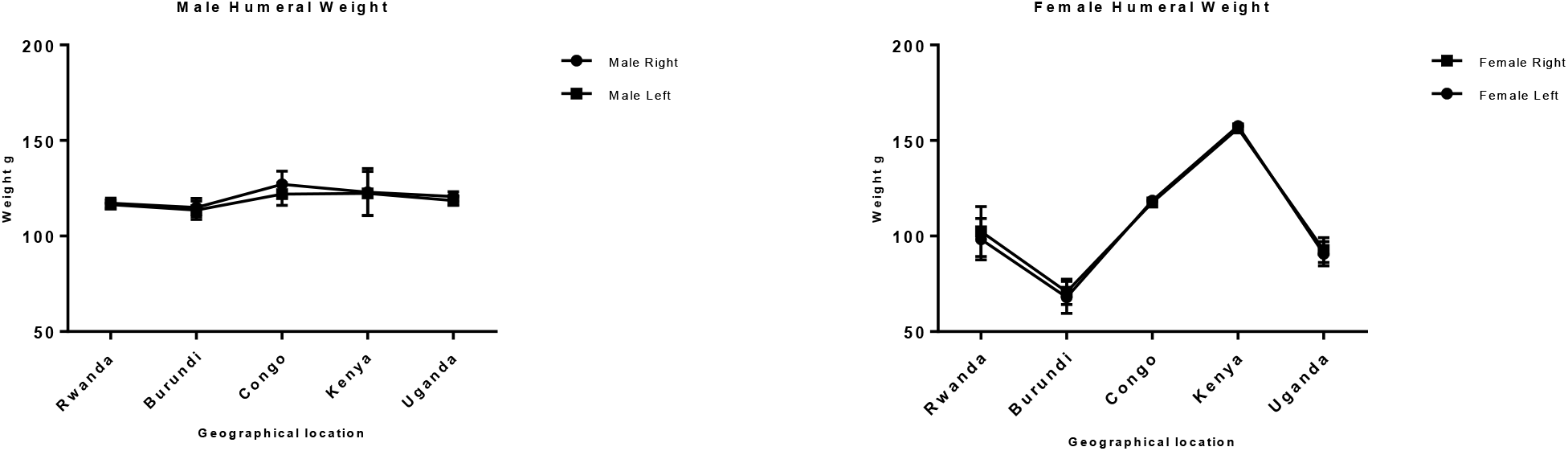
Shows the male and female right and left humeral weight across the geographical locations. (A) The right humerus is heavier than the corresponding left in all, correlation coefficient r = 0.94 and p = 0.019. (B) The right humerus is slightly heavier than the left in Rwanda, Burundi and Uganda while the left humerus is slightly heavier than the right in Congo and Kenya, r = 0.99 and p = 0.00004. The error bars represent the standard deviation of measurements for number of specimen measured (n) per geographical location.

**Figure 6:**
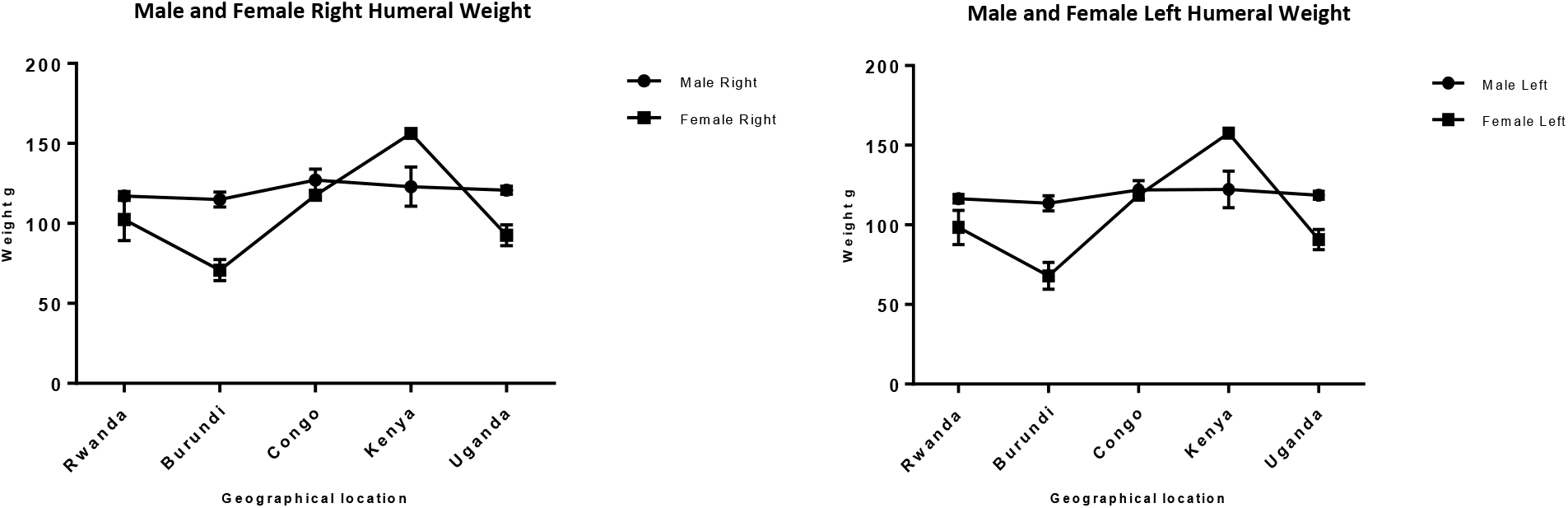
Shows a comparison between male and female humeral weight across the geographical regions. (A) The right male humerus is slightly higher in all except in Kenya where the right female is slightly higher than the male, r = 0.66 and p = 0.22 (B) The left male humerus is slightly heavier in all except in Kenya where the left female is slightly higher than the male, r = 0.88 and p = 0.04. The error bars represent the standard deviation of measurements for number of specimen measured (n) per geographical location.

Measurement of humeral length revealed longer right humeri in all geographical locations in both male and female except in Kenya where both male and female are of equal length. Comparison of male and female corresponding right and left shows a longer humerus in the male in all geographical location except in Kenya which is vice versa (Fig. 7&8). This observation in humeral length is in agreement with the report of Cuk et al., (2001) and Barros & Soligo (2013) which showed that average lateral asymmetry in the arms was to the right and that humans are unique in being lateralized to the right. Looking at the measurement of the mid-shaft circumference (Fig. 9&10), we also observed higher values on the right humeri in all the geographical locations except in Congo and Kenya female which have equal values on both right and left humeri (Table 1&2). These observations are similar to that of humeral weight with similar disparities. Furthermore, measurement of humeral torsion angle revealed an interesting phenomenon where angle of torsion is greater on all male left humeri except in Congo having a greater torsion angle on the right. The female specimen presents greater torsion angle on the left in all except Rwanda where angle of torsion in greater on the right. Burundi, Kenya and Uganda showed a consistent greater torsion on the left humerus compared to the right in both sexes. On the contrary Rwanda shows right greater torsion in male while Congo shows greater left torsion in female (Fig. 11&12). Our study therefore contradicted the report of Barros and Soligo (2013) which stated that both the magnitude and direction of asymmetries in humeral torsion in paired humeri from humans are unique in being lateralized to the right. However, in all the parameters measured in this study, we observed that right humerus in both sexes in most cases presented larger values compared with the left except the angle of torsion which is variable. This right-sided asymmetry may result from the normal tendency of individuals to favour the right upper limb during power activities (Kewal Krishan 2011).

**Table 1:**
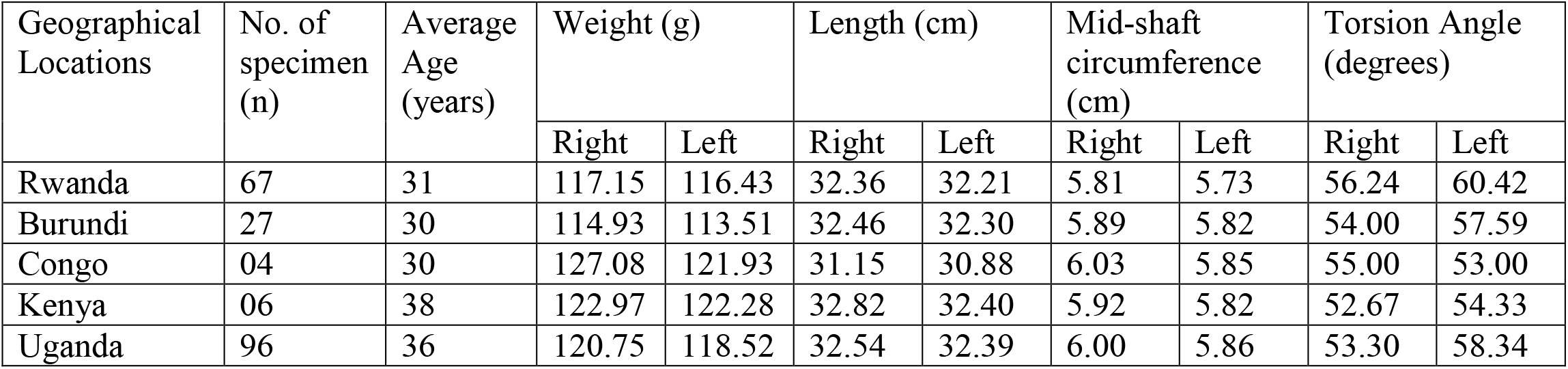
Shows male average age, weight, length, mid-shaft circumference and torsion angle across the geographical locations

**Table 2:** Shows female average age, weight, length, mid-shaft circumference and torsion angle across the geographical locations

| Geographical Locations | No. of specimen (n) | Average Age (years) | Weight (g) |  | Length (cm) |  | Mid-shaft circumference (cm) |  | Torsion Angle (degrees) |  |
| --- | --- | --- | --- | --- | --- | --- | --- | --- | --- | --- |
|  |  |  | Right | Left | Right | Left | Right | Left | Right | Left |
| Rwanda | 10 | 29 | 102.39 | 98.35 | 31.01 | 30.68 | 5.66 | 5.60 | 57.00 | 55.40 |
| Burundi | 02 | 38 | 70.80 | 67.95 | 30.60 | 30.05 | 4.90 | 4.90 | 51.50 | 57.50 |
| Congo | 01 | 35 | 117.60 | 118.60 | 31.50 | 30.90 | 5.60 | 5.60 | 47.00 | 60.00 |
| Kenya | 01 | 30 | 156.30 | 157.60 | 35.00 | 35.00 | 6.50 | 6.50 | 55.00 | 59.00 |
| Uganda | 20 | 32 | 92.64 | 90.72 | 30.59 | 30.20 | 5.49 | 5.45 | 55.70 | 58.20 |

**Figure 7:**
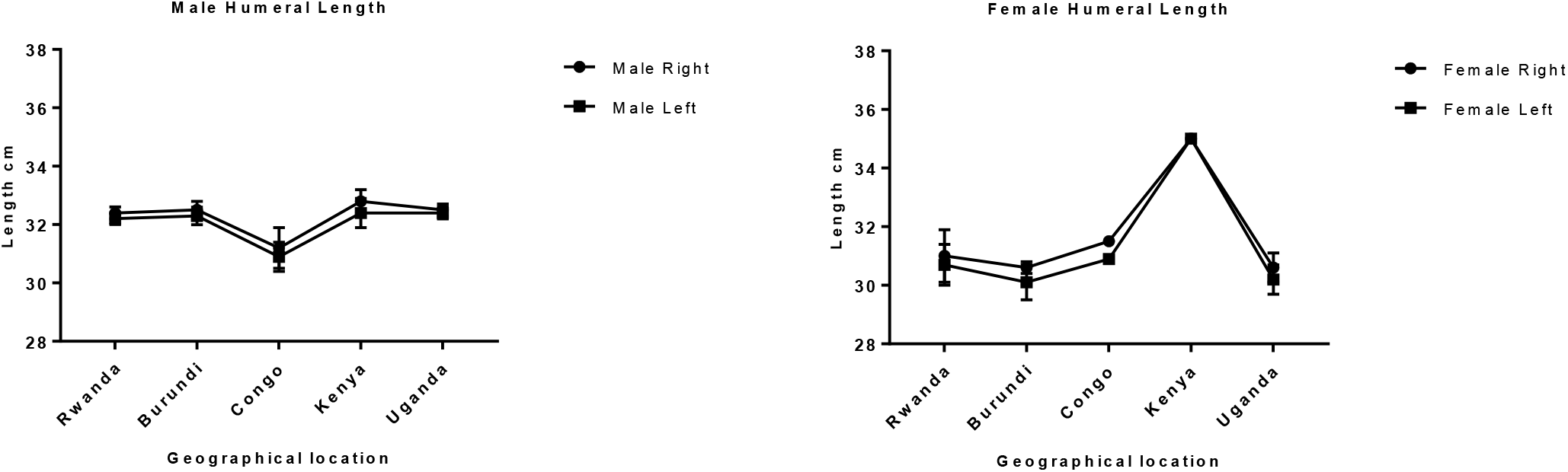
Shows the male and female right and left humeral length across the geographical locations. (A) The right male humerus is longer than the corresponding left in all the geographical locations, r = 0.98 and p = 0.002. (B) The right female humerus is longer than the left in all except in Kenya where the right is equal to left, r = 0.99 and p = 0.0001. The error bars represent the standard deviation of measurements for number of specimen measured (n) per geographical location.

**Figure 8:**
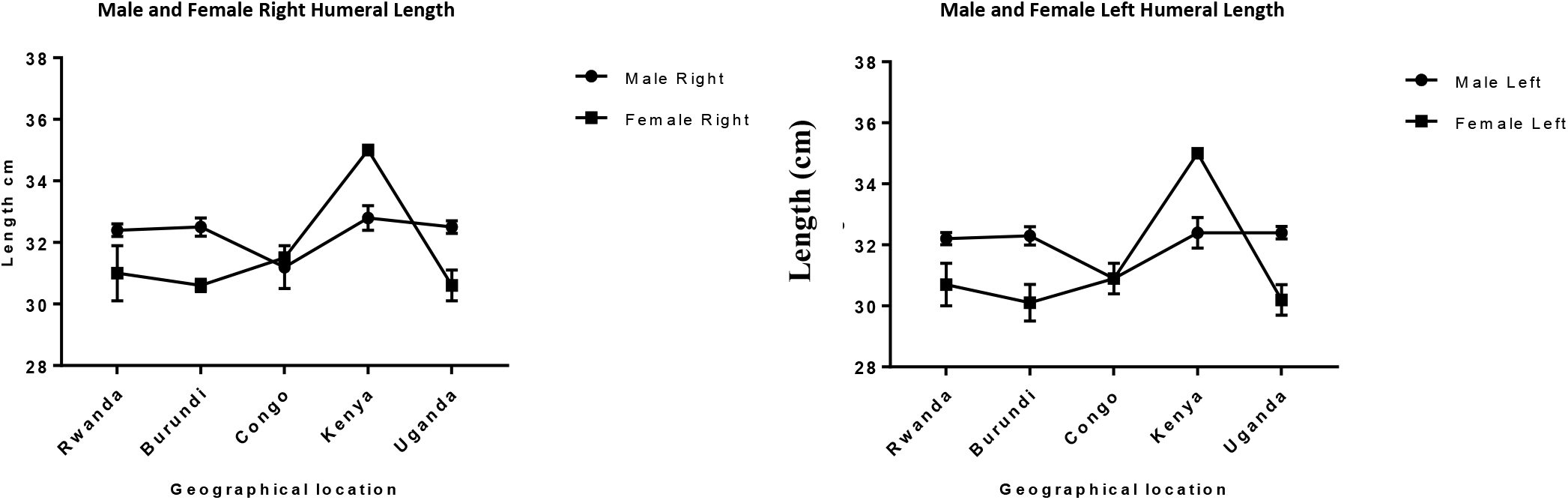
Shows a comparison between male and female humeral length across the geographical regions. (A) The right male humerus is longer in all except in Kenya where the right female is longer than the male, r = 0.3 and p = 0.63. (B) The left male humerus is longer in all except in Kenya where the left female is longer than the male, r = 0.19 and p = 0.76. The error bars represent the standard deviation of measurements for number of specimen measured (n) per geographical location.

**Figure 9:**
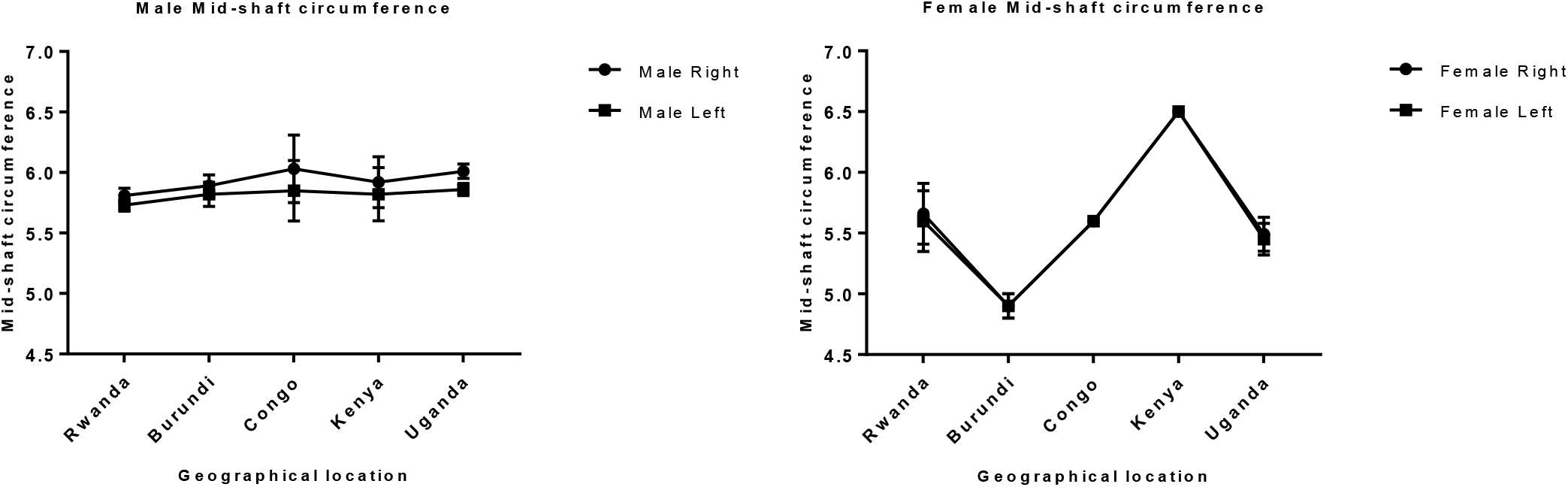
This chart shows the male and female right and left mid-shaft circumference of the humerus across the geographical locations. (A) The right male humerus is thicker than the corresponding left in all the geographical locations, r = 0.92 and p = 0.026. (B) The right female humerus is thicker than the left in all except in Kenya where the right is equal to left, r = 0.99 and p = 0.00005. The error bars represent the standard deviation of measurements for number of specimen measured (n) per geographical location.

**Figure 10:**
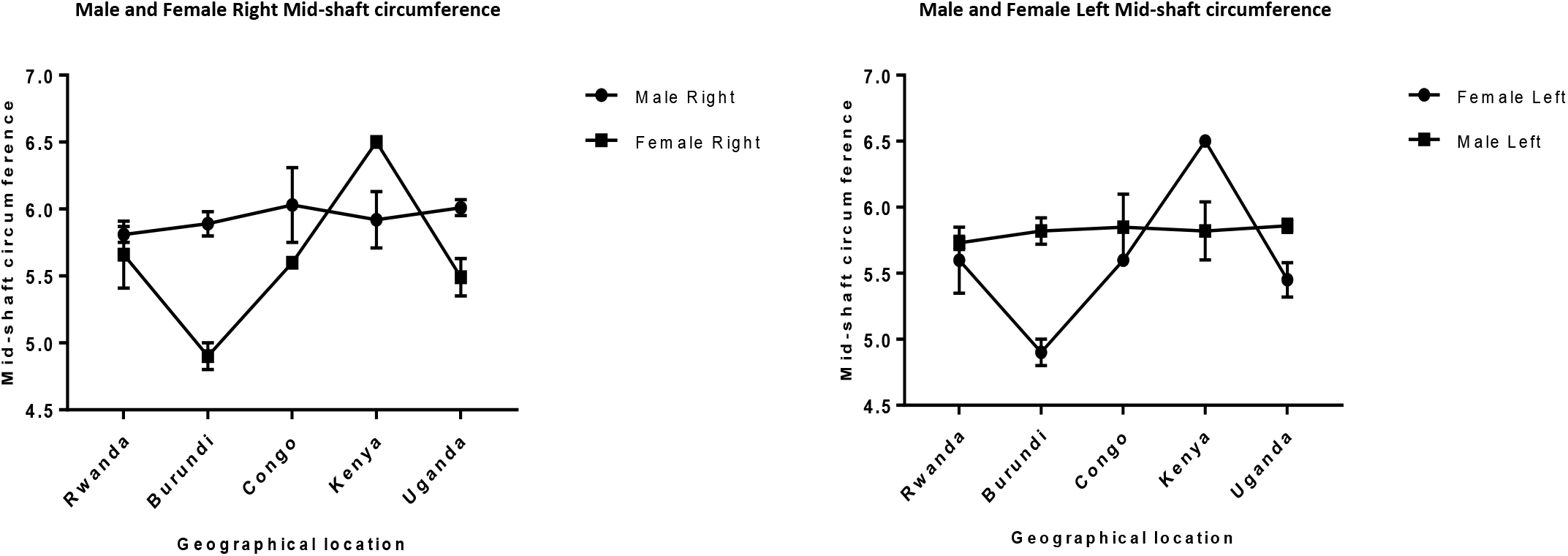
Shows a comparison between male and female mid-shaft circumference of the humerus across the geographical regions. (A) The right male humerus is slightly thicker in all except in Kenya where the right female is slightly thicker than the male, r = 0.013 and p = 0.98. (B) The left male humerus is slightly thicker in all except in Kenya where the left female is slightly thicker than the male, r = −0.049 and p = 0.94. The error bars represent the standard deviation of measurements for number of specimen measured (n) per geographical location.

**Figure 11:**
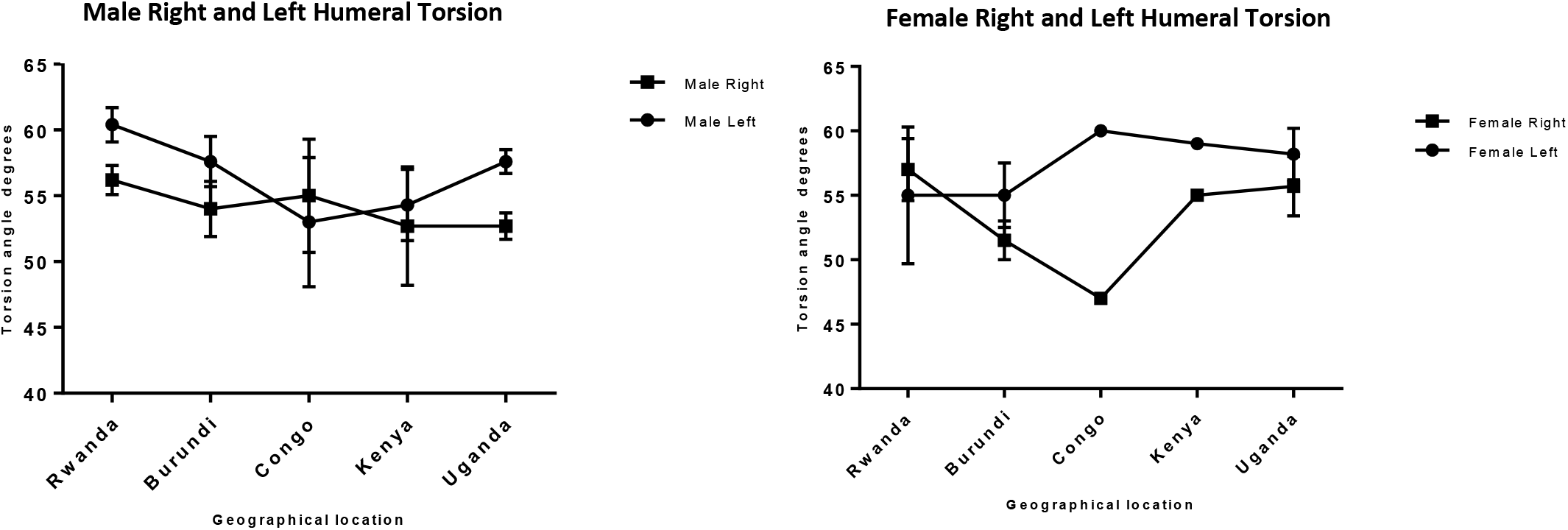
Shows the male and female right and left humeral torsion across the geographical locations. (A) The right male humerus is lesser than the corresponding left in both sexes across all the geographical locations except in Congo where the right is more than the left, r = 0.36 and p = 0.55. (B) The right female humerus is lesser than the left in all except in Rwanda where the right is more than the left, r = 0.44 and p = 0.46. The error bars represent the standard deviation of measurements for number of specimen measured (n) per geographical location.

**Figure 12:**
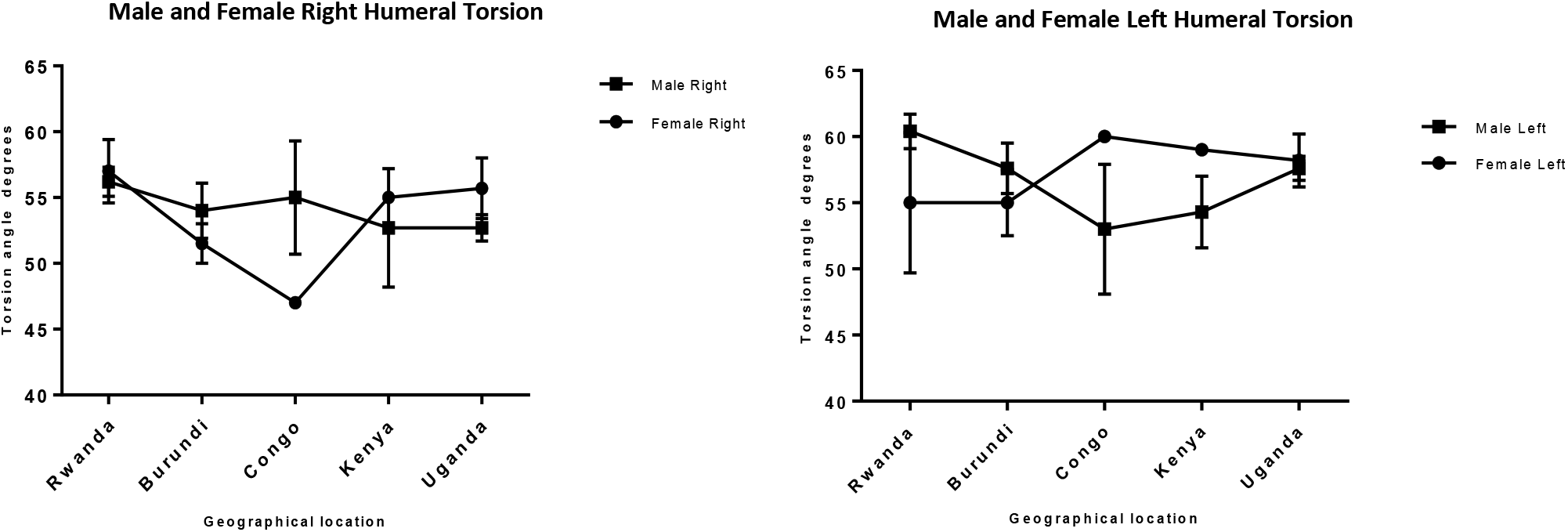
Comparison between male and female right and left humeral torsion across the geographical regions. (A) The right female torsion angle is more than male in Rwanda, Kenya and Uganda while the right male torsion is more than female in Burundi and Congo, r = −0.14 and p = 0.82. (B) The left female torsion is more than males in Congo, Kenya and Uganda while the left male torsion is more than female in Rwanda and Burundi, r = −0.87 and p = 0.055. The error bars represent the standard deviation of measurements for number of specimen measured (n) per geographical location.

According to our result, there is a strong correlation between the right and left humeral weight, length and mid-shaft circumference, but a slightly weak correlation in the torsion angle across the geographical locations (Fig. 5, 7, 9 & 11). Comparison between right and left of both sexes shows a strong correlation in humeral weight (Fig. 6), weak correlation in humeral length, mid-shaft circumference in male (Fig. 8 & 10) and no correlation in female mid-shaft circumference as well as humeral torsion angle in both sexes (Fig. 10 & 12).

According to Steel and Mays (1995) “The proportion of people with longer right or left long arm bones in the medieval population agreed with the proportion of right or left handers in modern population”. It was established that the main cause of oriented asymmetry is right or left handedness in that the arm subjected to greater mechanical stress as a result of use becomes the dominant. Generally, without a particular pathological condition, differential loading is most often a reflection of limb dominance (handedness) (Roy *et al.* 1994).

Findings have established that ipsilateral, as well as contralateral, movements activate the left, but not the right, motor cortex or associated areas of either hemisphere. Investigation of patients with damaged brain shows that the left hemisphere in right handers is specialized for controlling cognitive motor tasks in both arms. In the light of this future studies is needed to investigate the mechanisms for this asymmetry and to possibly establish a relationship between humeral asymmetry and anatomical characteristics of the brain.

## CONCLUSION

This study established the existence of bilateral asymmetries in the humerus of all the geographical regions investigated with more asymmetry observed in the male compared with the female. The asymmetry is to the right in the weight, length and mid-shaft circumference while the asymmetry is to the left in torsion angle. Also, our investigation revealed that humeral torsion is inversely proportional to weight, length and mid-shaft circumference of the humerus.

## DATA AVAILABILITY

The metadata used to support the findings of this study are available from the corresponding author upon request.

## AKNOWLEDGEMENTS

The researcher wish to acknowledge Department of Human Anatomy, Kampala International University Western Campus and Department of Human Anatomy School of Health Sciences Makarere University for giving access to the bones in the Osteological Collection Museum.

